# Saak Transform-Based Machine Learning for Light-Sheet Imaging of Cardiac Trabeculation

**DOI:** 10.1101/793182

**Authors:** Yichen Ding, Varun Gudapati, Ruiyuan Lin, Yanan Fei, Sibo Song, Chih-Chiang Chang, Kyung In Baek, Zhaoqiang Wang, Mehrdad Roustaei, Dengfeng Kuang, C.-C. Jay Kuo, Tzung K. Hsiai

## Abstract

Recent advances in light-sheet fluorescence microscopy (LSFM) enable 3-dimensional (3-D) imaging of cardiac architecture and mechanics *in toto*. However, segmentation of the cardiac trabecular network to quantify cardiac injury remains a challenge. We hereby employed “subspace approximation with augmented kernels (Saak) transform” for accurate and efficient quantification of the light-sheet image stacks following chemotherapy-treatment. We established a machine learning framework with augmented kernels based on the Karhunen-Loeve Transform (KLT) to preserve linearity and reversibility of rectification. The Saak transform-based machine learning enhances computational efficiency and obviates iterative optimization of cost function needed for neural networks, minimizing the number of training data sets to three 2-D slices for segmentation in our scenario. The integration of forward and inverse Saak transforms serves as a light-weight module to filter adversarial perturbations and reconstruct estimated images, salvaging robustness of existing classification methods. The accuracy and robustness of the Saak transform are evident following the tests of dice similarity coefficients and various adversary perturbation algorithms, respectively. The addition of edge detection further allows for quantifying the surface area to volume ratio (SVR) of the myocardium in response to chemotherapy-induced cardiac remodeling. The combination of Saak transform, random forest, and edge detection augments segmentation efficiency by 20-fold as compared to manual processing; thus, establishing a robust framework for post light-sheet imaging processing, creating a data-driven machine learning for 3-D quantification of cardiac ultra-structure.

## I. INTRODUCTION

Light-sheet fluorescence microscopy (LSFM) is instrumental in advancing the field of developmental biology and tissue regeneration [1-4]. LSFM systems have the capacity to investigate cardiac ultra-structure and function [5-14], providing the moving boundary conditions for computational fluid dynamics [15, 16] and the specific labeling of trabecular network for interactive virtual reality [17, 18]. However, efficient and robust structural segmentation of cardiac trabeculation remains a post-imaging challenge [19-23]. Manual segmentation of cardiac images in zebrafish and mice remains as the gold standard for ground truth despite being a labor-intensive and error-prone method. Currently existing automatic methods, including adaptive binarization, clustering, voronoi-based segmentation, and watershed, have remained limited [24]. For instance, adaptive histogram thresholding provides a semi-automated computational approach to perform image segmentation; however, the output is detracted by variability in background-to-noise ratio from the standard image processing algorithms [23].

Machine learning strategies, including neural networks [25-33], have been an integral part of biomedical research [34, 35] and clinical medicine [36-38]. The implementation of regions with convolutional neural network (R-CNN) provides the foundation for object detection and segmentation [29]. The fully convolutional network (FCN) also enables accurate and efficient image segmentation, replacing the fully connected layers with deconvolution-based upsampling and shared convolution computation [30]. The deep neural network (DNN) further allows for boundary extraction based on a pixel classifier in the electron microscopy images [26], and similar advanced neural networks facilitate the classification and segmentation of MRI images [28, 32].

However, these aforementioned methods require a large volume of high-quality annotated-training data, which is not readily available in the vast majority of LSFM-acquired cardiac trabecular network. Despite being an effective deduction method, a convolutional neural network (CNN) is considered to be a weak inference method to capture semantic information. In short, the limited resources of well-established ground truth and the weak inference capability motivated us to develop the subspace approximation with augmented kernels (Saak) transform, whereby we integrated the Karhunen-Loeve Transform (KLT) with the random forest classifier to reduce the need for high volume of annotated LSFM training data. We established a Saak transform to preserve linearity and reversibility of rectified linear unit (ReLU), and we applied Saak transform to extract the transform kernels for high energy spectral representations [39]. We paired KLT and augmented kernels with a random forest classifier to leverage the data efficiently to perform feature extraction and classification for LSFM image segmentation. We further integrated our Saak transform-based machine learning with LSFM imaging and edge detection of structural change in myocardium to define the surface area to volume ratio (SVR) in response to chemotherapy-induced injury. In addition, both forward and inverse transforms available in our Saak transform method enabled us to filter out adversarial perturbations and to reconstruct estimated images with guaranteed fidelity, for reducing a threat prone to errors in the neural networks [40-44]. Overall, Saak transform is a high throughput method to demonstrate precise and robust image segmentation of the complicated cardiac trabecular network, implicating both deductive and inference capacities to perform segmentation following cardiac injury.

## II. METHODS

### A. Fundamentals of Saak Transform

The overall Saak transform-based machine learning incorporates KLT kernels and augmented kernels for feature extraction and random forest for classification to generate a segmented image (Figure 1). In brief, we defined ***f***∈ *R*^*N*^ as the input, where *N* is the product of horizontal by vertical by spectral dimensions. The KLT basis functions are denoted by **b**_**k**_, *k = 1, 2*,…, *N* (Figure 1I, standard output of KLT), satisfying the orthonormal condition as follows:

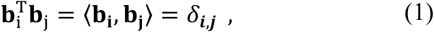

where *0* < *i, j ⩽ N* and *δ*_*i,j*_ is the Kroneckor delta function. A general approach after convolution in neural networks is to rectify the negative output to activate the non-linearity. However, we generated augmented kernels to preserve all the convolutional output, that is, the linearity of the transformation.

These augmented kernels enable us to comprehensively exploit image patterns for the feature extraction, defined as follows (Figure 1I, augmented output of KLT):

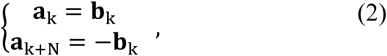

where **a**_**k**_ is the original KLT kernel while **a**_**k+N**_ is the augmented transform kernel. The integration of **a**_**k**_ and **a**_**k+N**_ allows for maintaining all the outputs during ReLu by assigning different signs. The input, ***f***, is then projected onto the augmented basis to yield a projection vector **p**, followed by ReLU and max pooling, and defined as follows:

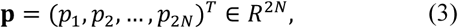

where 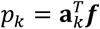. Since **a**_**k+N**_ is opposite to **a**_**k**_, we are able to convert the position to sign by on the projection vector:

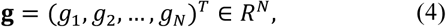

where *g*_*k*_ is a Saak coefficient, defined as follows:

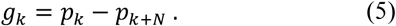

**Fig 1.**
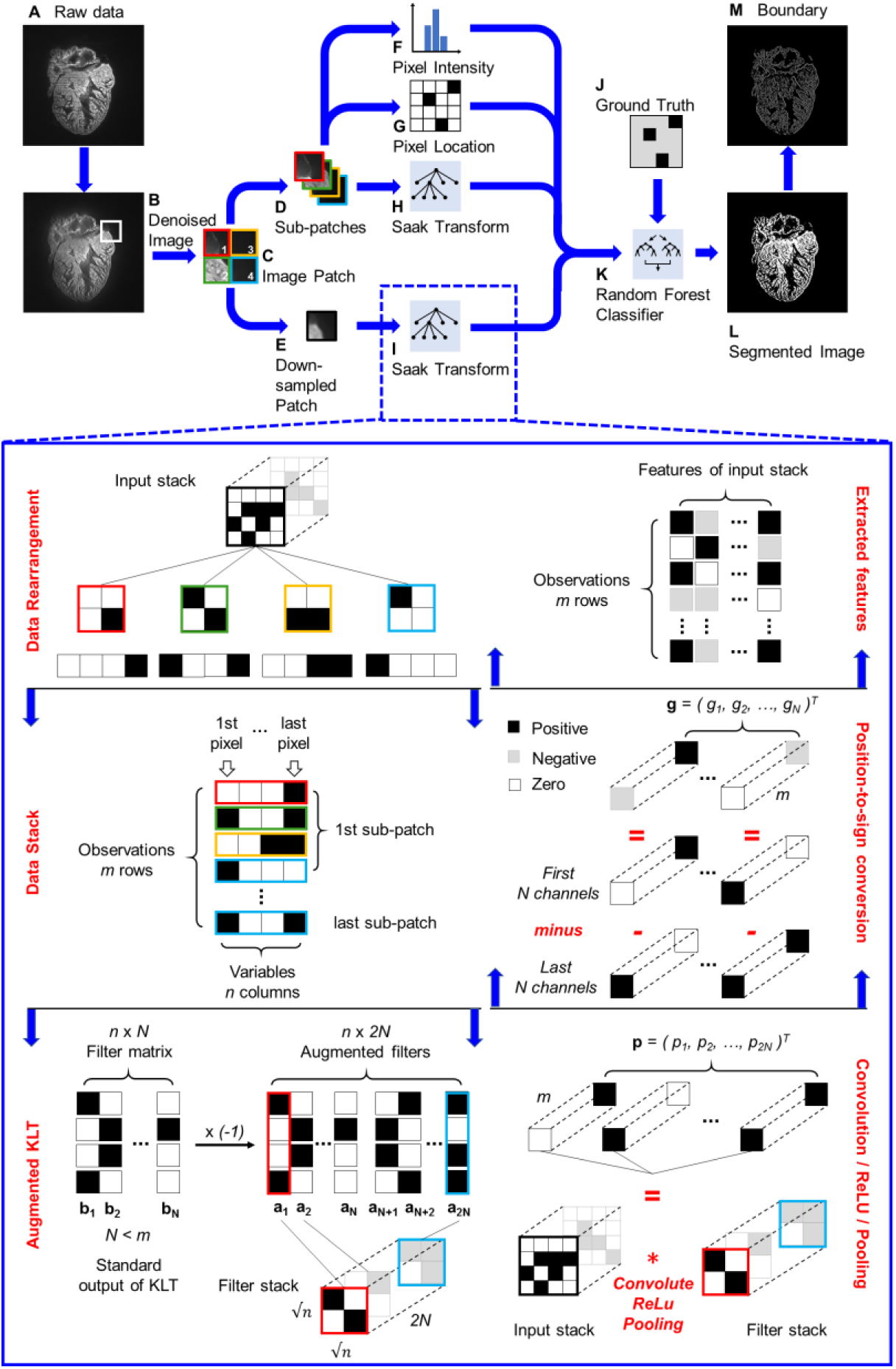
Pipeline of Saak transform-based machine learning. (A) The raw image is denoised by (B) variational stationary noise remover, and (C) split into patches. Patches are further reduced into (D) sub-patches and (E) down-sampled patches. (H and I) These two types of patches are passed to the Saak transform, which operates by performing an augmented Karhunen-Loeve Transform (KLT) operation, followed by the convolution, rectified linear unit (ReLU) and max pooling in a quad-tree structure. The feature extraction results of the KLT transform are packaged with (F) the pixel intensity and (G) location to train (K) a random forest classifier with (J) ground truth segmentation results. (L) The trained end-to-end algorithm has the capacity to segment image automatically, and (M) the segmented images are further processed by edge detection.

Thus, we finished a forward Saak transform by projecting the input, ***f***, onto transform kernels, **a**_**k**_ and **a**_**k+N**_, to generate Saak coefficients, *g*_*k*_, *k = 1,2*,…, *N*, as features of the input. Conversely, we are also able to reconstruct the estimation 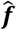 of input ***f*** after collecting Saak coefficients and their sign-to-position conversion:

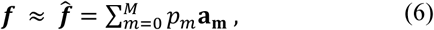

where *M* ≤ *2N*. This process is known as the inverse Saak transform, and we have provided mathematical analysis of Saak transform in details [39].

### B. Non-overlapping Saak Transform

We pre-processed raw LSFM images by using variational stationary noise remover (VSNR) to minimize the striping artifacts (Figures 1A, B) [45, 46].

Following the application of the denoising algorithm, the images of 1024 × 1024 pixels were decomposed into non-overlapping patches with 16 × 16 pixels (Figure 1C). An averaging window with 4 × 4 pixels further divided these patches into sub-patches (Figure 1D) and down-sampled patches (Figure 1E), respectively. Non-overlapping Saak transform operates on a quad-tree structure that splits the input into quadrants until reaching leaf nodes with 2 × 2 pixel squares (Inset of Figure 1I). The dimensions of the patch and sub-patches are application-dependent and adjustable, and the parameters were experimentally optimized. We used the following 4 parameters for feature extraction: pixel intensity (Figure 1F), pixel location (Figure 1G), Saak coefficients for sub-patches (Figure 1H), and Saak coefficients for down-sampled patches (Figure 1I). Our algorithm passed these four parameters to a random forest classifier (Figure 1K) for training with annotated data (Figure 1J). The algorithm also provides the optionality to select any classifier based on the application. The combination of feature extraction and classification created an end-to-end machine learning algorithm. Following the training, unannotated images were passed to generate a segmented result (Figure 1L) and a boundary image (Figure 1M).

All of the patches and sub-patches were passed to Saak transform for feature extraction. The input of Saak transform was generally a 3-D image stack such as a sub-patch or down-sampled patch stack. We first extracted blocks by sequentially walking the 2 × 2 window over the 2-D patches, and reshaped the block to 1-D array (Figure 1I, Data rearrangement). All the arrays were stacked along the column as a new 2-D matrix in which rows represent observations while columns are variables (Figure 1I, Data stack). We performed KLT on the matrix to extract the transform kernels **b**_**k**_, *k = 1, 2*,…, *N*, and reversed the sign of kernels to generate augmented kernels **a**_**k**_ and **a**_**k+N**_ (Figure 1I, Augmented KLT). All the kernels were reshaped back to 2 × 2 pixel squares as filters to convolute with the input stack, followed by ReLU and max pooling (Figure 1I, Convolution / ReLU / Pooling). We acquired coefficients, *p*_*1*_, *p*_*2*_,…, *p*_*2N*_, and subtracted *p*_*N+1*_, …, *p*_*2N*_ from *p*_*1*_, …, *p*_*N*_ to generate Saak coefficients, *g*_*1*_, …, g_*N*_, correspondingly. These coefficients enable to reduce the dimension of the final output and convert the position to sign, that is, the positive sign means the coefficient deriving from **a**_**k**_ while negative is from **a**_**k+N**_ (Figure 1I, Position-to-sign conversion). We rearranged all coefficients to finalize the features as the input of the classifier (Figure 1I, Extracted features).

An energy threshold between 0 and 1 (selected as 0.97 for this experiment) maintained the minimum accuracy of representation while truncating terms from the Saak representation for runtime efficiency. The accuracy of the Saak representation was verified with the analysis of variance (ANOVA) to complete the feature extraction process.

### C. Overlapping Saak Transform

In addition to the non-overlapping Saak transform, we generated overlapping patches for feature extraction. The main difference between two methods is the overlapping area among different patches. Following denoising, the Saak algorithm decomposed the image of 1024 × 1024 pixels into overlapping patches with 27 × 27 pixels. A 9 × 9 pixel averaging filter divided patches into sub-patches and down-sampled patches, respectively. All of the patches and sub-patches were passed to the Saak transform operating on a quad-tree structure that split the input into quadrants until reaching leaf nodes with 3 × 3 pixel squares. The dimensions of the patch and sub-patches are adjustable for application-dependent tasks.

### D. Light-Sheet Microscope Design

Our custom-built LSFM system was designed to image the changes in trabecular network in the zebrafish model of cardiac injury and regeneration. Our LSFM system adapting from [23, 47] generated a Gaussian light-sheet at ∼4.5 *µm* in thickness (Figure 2), and the detection unit was perpendicularly positioned to the illumination plane. Fluorescence was orthogonally captured through an objective lens (Ob1) and a tube lens (TL1) to minimize photobleaching while providing high axial resolution (Inset in Figure 2A). A sCMOS (Flash 4.0, Hamamatsu, Japan) was used to capture 1024 × 1024 pixels per image at 100 frames per second. Following precise alignment with the LSFM system, a fully automated image acquisition was implemented via a customized LabVIEW program. We created a virtual spiral phase plate (SPP) for automatic edge detection by performing the forward and inverse Fourier transform in MATLAB (MathWorks, Inc.) [48, 49]. The hardware system is also adaptable for inserting a real SPP along with optical components (Ob2 and TL2) as a high-pass filter for edge detection. For the sake of Saak transform, we chose to perform the edge detection and trim the boundary of cardiac trabeculae (Figure 2C) in MATLAB following LSFM imaging (Figure 2B).

**Fig 2.**
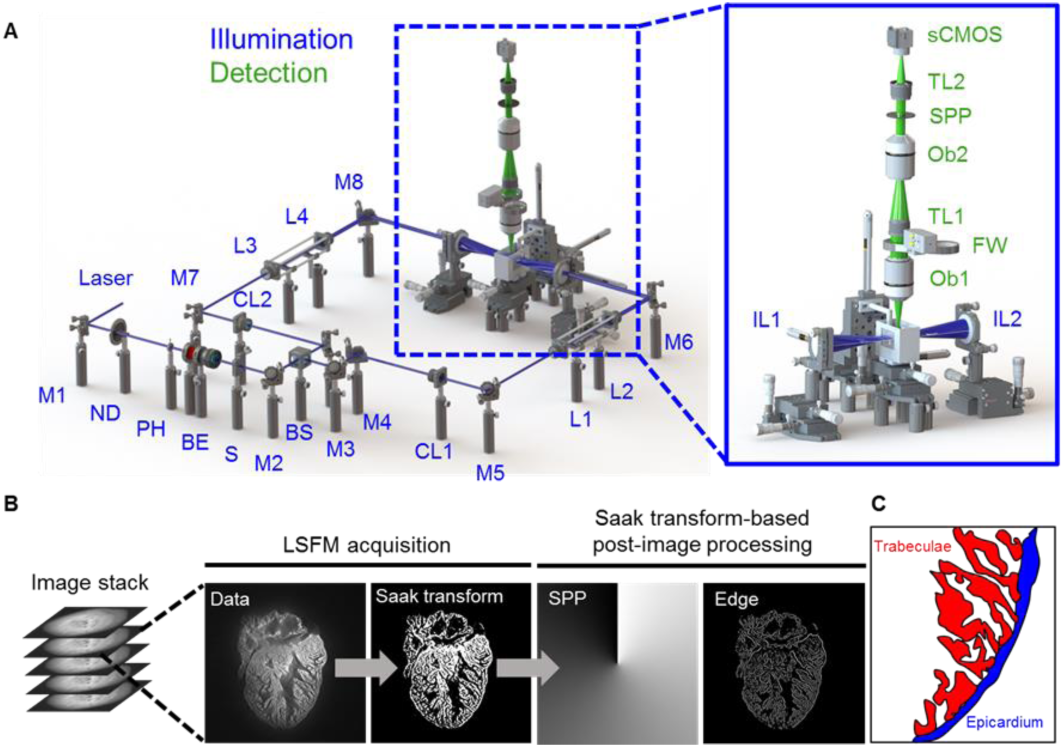
LSFM data acquisition and post image processing. (A) In the illumination unit (blue), the laser is reflected by a mirror (M) to a neutral density filter (ND), adjustable pinhole (PH), beam expander (BE), and slit (S). The beam is 1) split by a beam splitter (BS), 2) converted to a light sheet by the cylindrical lens (CL), and 3) tuned by relay lenses (L). Illumination lenses (IL) project the light-sheet across the sample. The detection unit (green) consists of two objective lenses (Ob) and two tube lenses (TL) to focus the image on the sCMOS camera. The filter wheel (FW) and optional spiral phase plate (SPP) are used to further process the image. (B) The raw image stack acquired by LSFM is segmented through Saak transform-based machine learning, followed by the edge detection and boundary trim. (C) A schematic highlights the cardiac trabecular network (red) in the adult zebrafish ventricle (blue).

### E. Generation of Adversarial Perturbation

We implemented well-established perturbation methods, including Deepfool [50], fast gradient sign (FGS) [51] and gradient ascent (GA) methods [52], to generate adversarial perturbed images. Adversarial perturbation is designed to challenge the robustness of classification by machine learning networks. We included both targeted and untargeted attacks in the FGS method [51]. In comparison to FGS and GA methods, Deepfool leads to a smaller perturbation. We used random forest as the classifier, and compared the segmentation output between the perturbed and reconstructed images.

### F. Saak Coefficients Filtering

Clean and adversarial images shared similar distributions in the low-frequency Saak coefficients despite variability in the high-frequency coefficients [44]. To reduce adversarial perturbations, we sought to truncate the high-frequency Saak coefficients as a filtering strategy, while the low-frequency Saak coefficients were maintained to reflect the continuous surface or slow varying contour in the segmental process. Following the truncation of high-frequency Saak coefficients generated by two-stage forward Saak transform, we used the rest of the coefficients to reconstruct the image based on the inverse Saak transform (Figure 5). Two types of moving windows were applied: 4 × 4, and 2 × 2 pixels. In the grayscale image, we truncated 200 (73% of total), 150 (55%), 100 (37%) and 50 (18%) coefficients by using a 4 × 4 moving window, and we also truncated 20 (95% of total), 15 (71%), 10 (48%) and 5 (24%) coefficients by using a 2 × 2 moving window. In addition, we performed the same procedure in the binary image, and filtered out 50 (18% of total), 40 (15%), 30 (11%) and 20 (7%) coefficients by using the 4 × 4 moving window, along with 20 (95% of total), 15 (71%), 10 (48%) and 5 (24%) coefficients by using the 2 × 2 moving window. All of the collected Saak coefficients were applied in inverse Saak transform to reconstruct the images.

**Fig 5.**
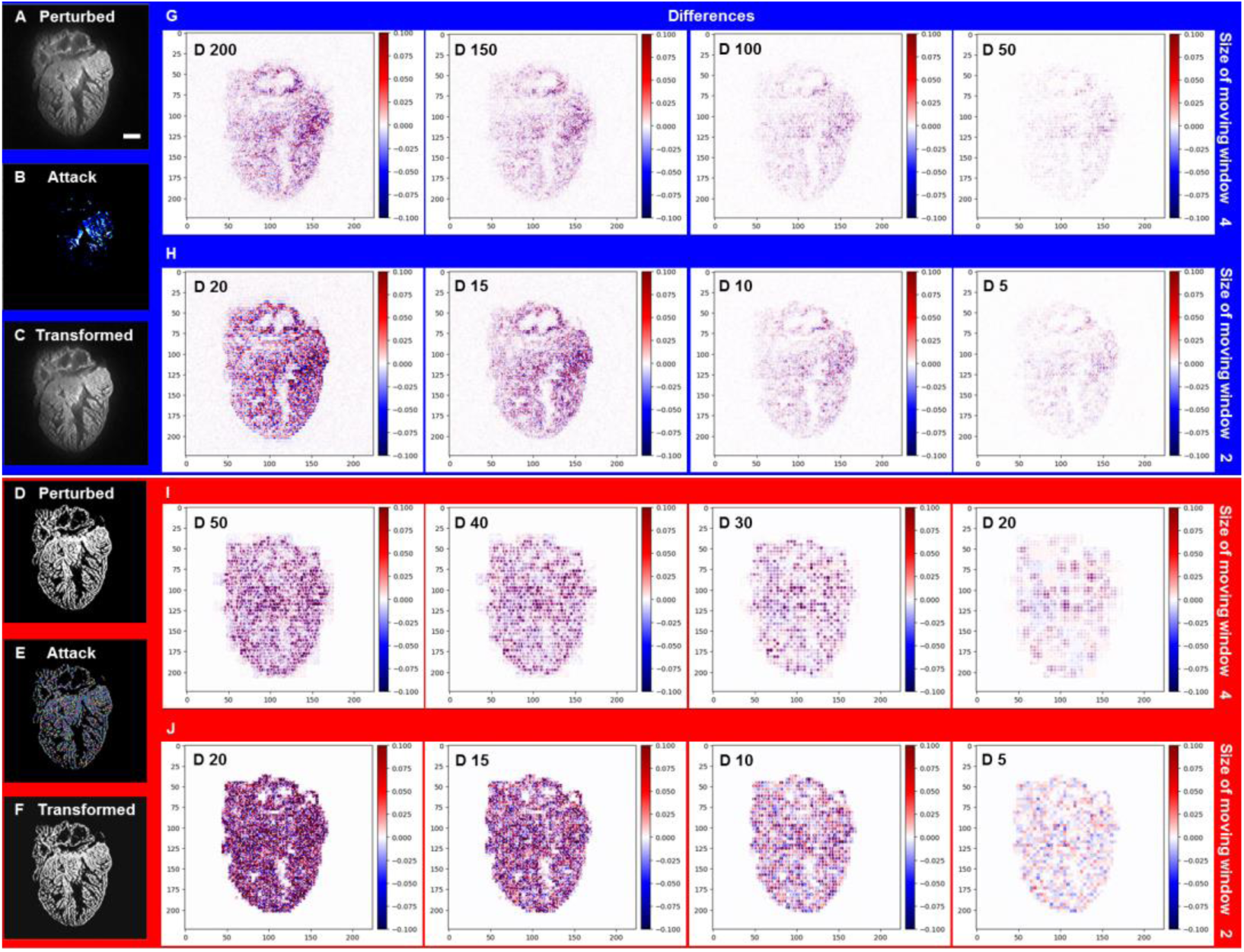
Forward and inverse Saak transforms to filter the adversarial perturbations from LSFM images. (A and D) Adversarial perturbed images were generated by Deepfool. (B and E) Perturbations were added to both grayscale and binary images. (C and F) Transformed images were reconstructed following truncation of the high-frequency coefficients. (G) By applying the 4 × 4 moving window, 200 (73% of total), 150 (55%), 100 (37%) and 50 (18%) coefficients (spectral dimensions) were filtered out from the grayscale image, respectively, and (I) 50 (18% of total), 40 (15%), 30 (11%) and 20 (7%) coefficients were filtered out from the binary image, respectively. (H and J) By applying the 2 × 2 moving window, 20 (95% of total), 15 (71%), 10 (48%) and 5 (24%) coefficients were filtered out from both the grayscale and binary images, yielding the representative differences from the binary image. The pseudo-color represents the intensity of specified adversarial perturbation. D stands for number of Saak coefficients. Scale bar: 200 *µm*.

### G. Assessing Changes in Cardiac Trabecular Network Following Chemotherapy-Induced Injury

All animal studies were performed in compliance with the IACUC protocol approved by the UCLA Office of Animal Research. Experiments were conducted in adult 3-month old zebrafish (*Danio rerio*). A one-time 5μL injection of doxorubicin (Sigma) at a dose of 20μg/g of body weight by intraperitoneal route (Nanofil 10μL syringe, 34 gauge beveled needle, World Precision Instruments) [53] or of a control vehicle (Hank’s Balanced Salt Solution) was performed at day 0 (n=5 for doxorubicin, and n=5 for control). Imaging of autofluorescence was performed using a custom-built LSFM system at day 30 post injection, followed by Murray’s clear to render translucency for imaging.

### H. Preparation of the Ground Truth

The image stack was acquired with a 2 *µm* step size yielding a resolution at 1.625 *µm* × 1.625 *µm* × 2 *µm* and an image size at 1024 × 1024 pixels. We manually segmented the myocardium with binary labels in Amira 6.1 (FEI, Berlin, Germany). Pixels that represented myocardium were flagged with a value of 1, while the cavities and spaces were flagged with a value of 0. The binary segmented images served as either training data for the Saak transform algorithm or validation sets to verify the accuracy of the Saak transform-based image segmentation.

### I. Validation

The performance of the segmentation algorithm was validated by calculating the dice similarity coefficient (DSC),

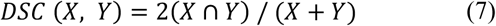

where *X* represents the ground truth segmented myocardial region, and *Y* the Saak transform or U-Net segmented myocardial region. The algorithm was tested with varying numbers of training patches.

### J. Statistical Analysis

All of values were represented as mean ± standard deviation. A two-sample *t*-test was used to assess the means between two data sets, followed by a two-sample *F* test for variance and a Kolmogorov-Smimov test for normality. All the comparable groups under each condition were subject to normal distribution and equal variance. In the box plot, the box was defined by the 25th and 75th percentiles, and the whiskers were defined by the 5th and 95th percentiles. Median, mean and outliers were indicated in line, dot, and diamond, respectively. A *p*-value of < 0.05 was considered statistically significant.

### K. Code availability

All of the Saak transform codes were written and tested in Python 3.6. The computer code generated during and/or analyzed during the current study are available from the corresponding author.

### L. Data availability

The data sets generated during and/or analyzed during the current study are available from the corresponding author.

## III. RESULTS

### A. Qualitative Comparison of the Segmented Trabecular Network

We compared the non-overlapping and overlapping Saak transforms with the well-recognized U-Net. We implemented both non-overlapping and overlapping patches to generate transform kernels and the corresponding Saak coefficients by training with 1, 3, 6, 9, 18 and 27 images (Figures 3A-B). Using the same number of training images (TI), Saak transform captured the trabecular network in greater detail compared to U-Net (Figure 3C). By comparing segmentation accuracy from 3 to 27 training images (Figures 3D-F), we qualitatively assessed the identical region of ventricle (dashed green squares in Figure 3G) to demonstrate that the Saak transform is superior with regards to segmentation of the interconnected trabecular network. We revealed the differences between the non-overlapping- and overlapping-patches Saak algorithms by comparing them with the manually annotated ground truth under the identical training conditions (Figure 3G). The non-overlapping-patches method was sensitive to high-resolution image components, revealing the detailed trabecular formation at the cost of noisy background, whereas the overlapping-patches method segmented a smooth and interconnected trabecular formation with lower resolution but less noise. We further highlighted the distinct results between the non-overlapping and overlapping methods by testing the Saak transform with a single training image (TI 1) (Figures 3H-I). With the reduced volume of training images needed to segment cardiac structure, Saak transform-based machine learning outperformed the well-established U-Net from a data-efficiency standpoint. However, the non-overlapping method yielded pixelated trabecular structures when only provided with 1 training image, whereas the overlapping method result captured more detail in the ventricular trabecular formation. For these reasons, we employed over 3 training images to offset the trade-off between segmentation accuracy and training efficiency in the ensuing results.

**Fig 3.**
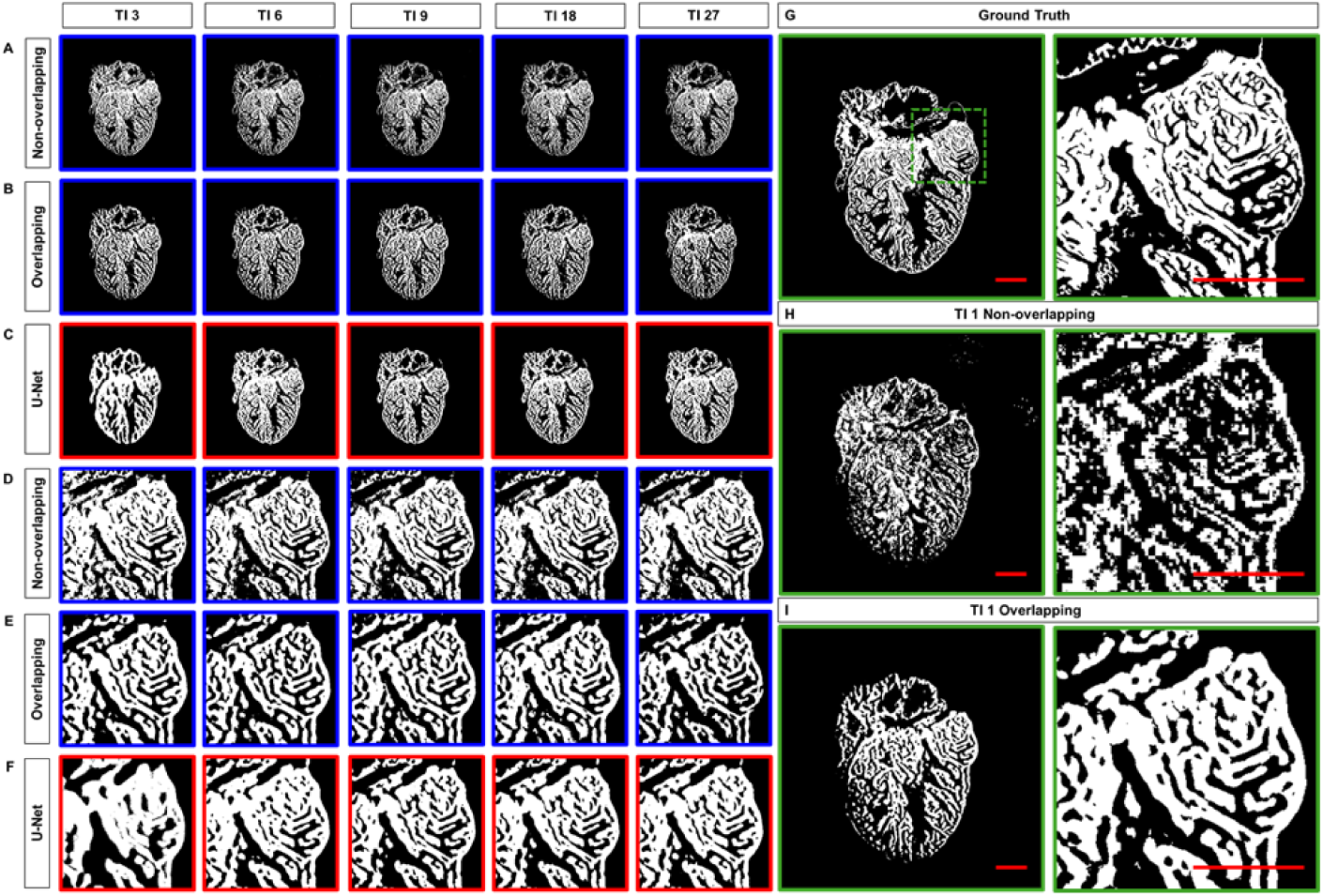
Comparison of Segmentation results. (A) Non-overlapping and (B) overlapping patches to generate Saak transform coefficients. (C) Under the same 3, 6, 9, 18 and 27 training images, the trabecular network is better demarcated by Saak transform than by U-Net. (D-F) Comparison of trabecular network in the region of interest (dashed square in G) further supports the capacity of Saak transform to reveal the detailed trabecular network. (H-I) The results of 1 training-based image are compared to the (G) ground truth. TI: training image. Scale bar: 200 *µm*.

### B. Quantitative Comparison of the Segmented Trabecular Network

We assessed the accuracy by comparing the correctly segmented regions of ventricle among 1) the non-overlapping Saak transform (Figure 4A, blue top channel), 2) overlapping Saak transform (Figure 4B, blue middle channel), and 3) U-Net (Figure 4C, red bottom channel) in relation to the manually annotated ground truth (Figures 4A-C, white merge). Dice similarity coefficient (DSC) was calculated to quantify the performance of segmentation following various numbers of training images. Based on 3 training images, we arrived at a mean DSC of 0.82 ± 0.07 for non-overlapping Saak transform and 0.72 ± 0.07 for U-Net (*p* = 0.001) (Figure 4D). The accuracy of segmentation following 3 training images for the non-overlapping method is comparable to that of 18 and 27 images for the same method; that is, the mean DSC value ranges from 0.80 to 0.90 for the non-overlapping Saak transform. Increasing the number of training images for U-Net improves the accuracy and approaches that of the non-overlapping Saak transform. Similar to the non-overlapping method, the overlapping Saak transform outperforms U-Net when provided with 3 training images, as demonstrated by a DSC value of 0.78 ± 0.04 (*p* = 0.02 for overlapping vs. U-Net). However, U-Net surpasses overlapping Saak transform, starting from 9 to 18 to 27 training images, yielding p-values from 0.001 to 0.003 to 0.0003, respectively. These findings corroborate the previous qualitative comparison (Figure 4A-C) between Saak transform-based machine learning and U-Net, supporting that the segmentation accuracy and training efficiency of Saak transform outperforms iterative optimization in neural networks.

**Fig 4.**
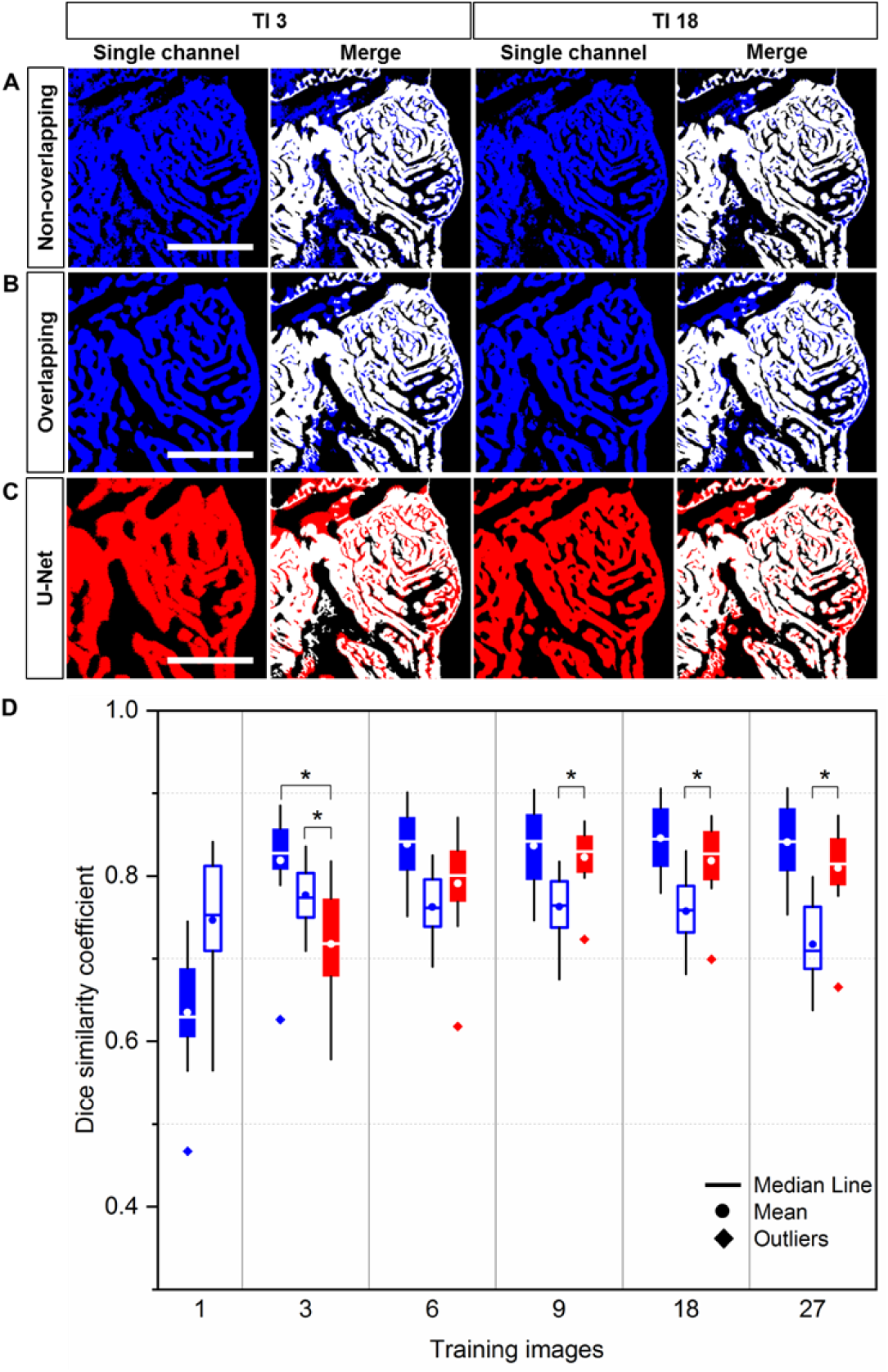
Quantitative comparison of Segmentation: Saak transform vs. U-Net. (A-C) Saak transform (in blue) vs. U-Net (in red) were compared following 3 and 18 training images. These segmentation results were merged with the manually annotated image (in white) for quantitative comparison. (D) Following 3 training images, the values of dice similarity coefficient (DSC) support that both non-overlapping (solid blue) and overlapping (hollow blue) Saak transforms outperformed U-Net (red). Starting from 9 to 18 to 27 training images, non-overlapping Saak transform and U-Net performed similarity, while overlapping Saak transform underperforms. Twelve testing patches were included in method under the identical training images. Solid blue: non-overlapping; hollow blue: overlapping; red: U-Net. Scale bar: 200 *µm*.

### C. Forward and Inverse Saak Transforms to Filter Adversarial Perturbations

In addition to reducing training images, the Saak transform allows for 1) filtering the high-frequency adversarial perturbation by using the forward Saak transform and 2) reconstructing the images with the inverse Saak transform. Using the well-established Deepfool attack, we added perturbation to both the grayscale (Figures 5A-C) and binary images (Figures 5D-F). Saak transform provides various moving windows, including 4 × 4 and 2 × 2 pixels, to extract coefficients to detect the perturbation deeply involved in the ventricular structure (Figures 5B and E). We truncated the high-frequency coefficients at different percentage as previously reported [44]. By applying the 4 × 4 moving window, we filtered out 200 (73% of total), 150 (55%), 100 (37%) and 50 (18%) coefficients from the grayscale image (Figure 5G), and 50 (18% of total), 40 (15%), 30 (11%) and 20 (7%) coefficients from the binary image (Figure 5I). By applying the 2 × 2 moving window, we repeated the procedure to reconstruct the image based on the remaining Saak coefficients (Figures 5H and J), including the case in which 95% of coefficients were removed (D stands for the number of coefficients: 20). Thus, both forward and inverse Saak transforms preserve robustness and enhance the authenticity of image reconstruction in the presence of adversarial perturbation.

### D. Saak Transform-based Segmentation of Perturbed and Reconstructed Images

To establish the robustness of image segmentation, we implemented Saak transform as a module to the random forest. Four images were perturbed by Deepfool, targeted FGS, untargeted FGS and GA methods, respectively (Figure 6A), and they were similar to that of the previous ground truth (Figure 1B). Next, adversarial noises were artificially added to these images (Figure 6B). We applied a two-stage non-overlapping Saak transform to segment both the perturbed (Figure 6C) and reconstructed images (Figure 6D), and we conducted a paired *t*-test following the individual perturbations (Figure 6E). We demonstrate that the low-frequency Saak coefficients are robust to multiple adversarial perturbations to the trabecular network, and the segmentation output reveals no statistically significant difference between the perturbed (blue) and reconstructed (red) segmentation (p = 0.95, 0.95, 0.96, and 0.96, twelve testing patches for each perturbation). In this context, Saak transform is a light-weight module for feature extraction in conjunction with the existing machine learning-based classifier.

**Fig 6.**
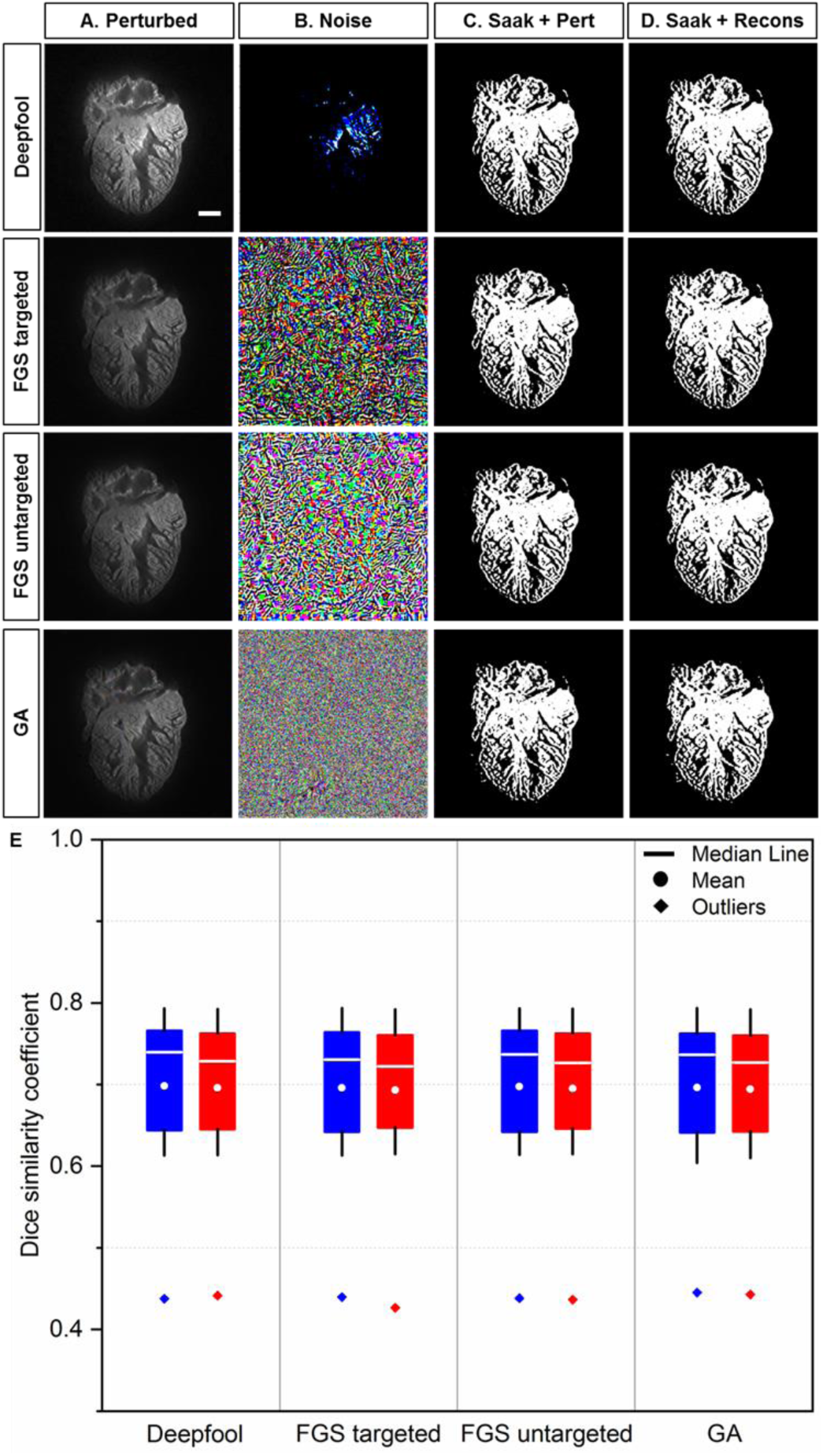
Saak transform-based segmentation following perturbed and reconstructed images. (A) Perturbed images are generated by introducing the original images to (B) adversarial perturbations from Deepfool, targeted and untargeted fast gradient sign (FGS), and gradient ascent (GA) methods, respectively. (C-D) Non-overlapping Saak transform plus random forest were performed to segment the cardiac trabecular network. (E) The values of DSC between perturbed and reconstructed images were statistically insignificant (*p* >0.05, n=12 testing patches for each perturbation). Blue: perturbed; red: reconstructed. Scale bar: 200 *µm*.

### E. Integrating Edge Detection to Assess Chemotherapy-Induced Changes in Trabeculation

To assess the chemotherapy-induced cardiac injury, we integrated Saak transform-based machine learning and edge detection for efficient segmentation. The Saak transform provides the fundamental basis to segment the 3-D endocardium and epicardium, enhancing the efficiency by 20-fold as compared to the manual segmentation. About 20 hours are required to segment a single 2-D image by a well-trained expert (Figure 7A), whereas only 1 hour is needed for Saak transform to complete both training and testing. In light of the complicated cardiac trabecular network in the adult zebrafish, we demonstrate the application of SVR as an indicator to assess the degree of trabecular network. We performed non-overlapping Saak transform in 5 pairs (doxorubicin treatment vs. control), and each treatment or control group contains 10 2-D images. The epicardium and endocardium were classified as white pixels, and the ventricular cavity was in black similar to the background. The white pixel counts were defined as the volume of cardiac structure (Figure 7A and C). We extracted the contour from the binary images, and trimmed the contour line with a single pixel filter (Figure 7B and D). The white pixels along the contour were defined as the surface area in 3-D. We summed all of the surface areas and volumes respectively, to compute SVR for the treatment vs. control groups. In response to chemo-induced cardiac injury, the SVR was significantly reduced (doxorubicin: SVR= 0.10 ± 0.02, n=5; control: SVR= 0.14 ± 0.03, n=5; *p* = 0.03). This finding was consistent with the previously reported reduction in cardiac trabeculation in response to chemo-induced myocardial injury [13, 23].

**Fig 7.**
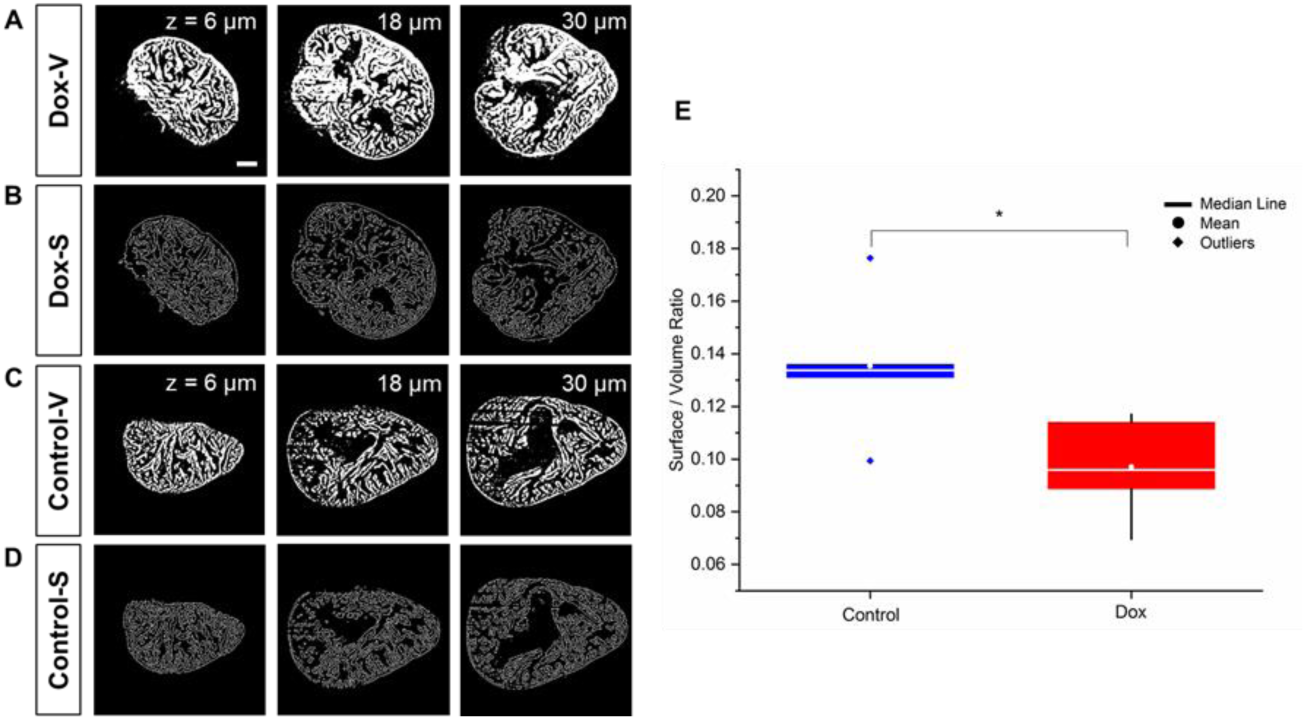
The surface-to-volume ratio (SVR) in response to doxorubicin treatment vs. control. (A) Saak transform-generated endo- and myocardial segmentation. (B) Surface contour provides a quantitative insight into cardiac remodeling in response to chemotherapy-induced heart failure. (C-D) The wild-type zebrafish were used as the control. Columns from left to right indicate different imaging depth at 6, 18, and 30 *µm*. (E) The reduction in SVR value is statistically significant following doxorubicin treatment (*p* = 0.03 vs. control, n=5 for treatment and control). Scale bar: 200 *µm*.

## IV. DISCUSSION

The advances in light-sheet imaging usher in the capacity for multi-scale cardiac imaging with high spatiotemporal resolution, providing a platform for the supervised machine learning methods to perform high-throughput image segmentation and computational analyses. However, manual segmentation of cardiac images is labor-intensive and error-prone. The main advantage of Saak transform-based machine learning is to reduce the demand for high-quality annotated data that are not readily available in most of LSFM imaging applications, thus allowing for a wide range of post-processing techniques in image segmentation. By partitioning the task into the feature extraction module and classification module, the Saak transform has the capacity to reduce the amount of annotated training data to segment the adult zebrafish hearts while remaining robust against perturbation.

The Saak transform-based machine learning encompasses 3 key steps: 1) to perform KLT on the input to build the optimal linear subspace approximation with orthonormal bases, 2) to augment each transform kernel with its negative, and 3) to apply the ReLU to the transform output. Unlike other CNN-based machine learning methods, the Saak transform for feature extraction is based on linear algebra and statistics and thus, provides mathematically interpretable results. Specifically, the Saak transform builds on the one pass feedforward manner to determine the coefficients through augmented KLT kernels, maximizing computational efficiency to obviate the need for determination and optimization of filter weights via training data and backpropagation. While a single training image is insufficient for U-Net to converge, the Saak transform-based machine learning method initiates image segmentation under the same conditions. We found that the Saak transform performs more accurate segmentations of LSFM structures than U-Net (at TI3, *p* = 0.001 for non-overlapping vs. U-Net, and *p* = 0.02 for overlapping vs. U-Net). The discriminant power of Saak coefficients is built on augmented kernels for pattern recognition and segmentation, and annotated images are only used in the kernel determination module instead of the feature extraction module. Thus, the number of training data sets decouples the segmentation accuracy, implicating that Saak transform has both deductive and inference capacities.

The Saak transform is robust against adversarial perturbations in input images if the covariance matrices are consistent. We have demonstrated that Saak transform-based machine learning outperformed other state-of-the-art methods in combating adversarial perturbations. Saak transform enables complement spatial smoothing techniques to mitigate the adversarial effects without interfering with the classification performance on raw images. In contrast to the unclear reversibility of CNN methods, the existence of the inverse Saak transform is well-defined. We are able to reconstruct the estimated images following the removal of high-frequency Saak coefficients that might be adversarial perturbed. The transparency of the Saak transform pipeline provides reproducibility to analyze breakdown points for numerous applications. All of the aforementioned features enable the Saak transform to dovetail for high throughput image segmentation, implicating an efficient and robust alternative to CNN.

The application of Saak transform to segment 3-D LSFM-acquired images creates an opportunity to investigate chemotherapy-induced cardiac structure and mechanics. Previously, we needed to perform manual or semi-automated segmentation to reconstruct LSFM architectures for computational fluid dynamics [15, 16] and interactive virtual reality [17, 18]. The addition of edge detection to Saak transform enables the calculation of surface area in relation to the volume of myocardium. The extent of trabecular network as measured by SVR was significantly attenuated following chemotherapy-induced myocardial injury. Conventionally, total heart volumes were measured by bounding the manual segmented results and filling the endocardial cavities, while endocardial cavity volumes were calculated by the difference between the total heart volumes and segmented myocardial volumes [23]. However, assessing endocardial cavity volume is a labor-intensive task. Based on the Saak transform and edge detection, we utilized SVR to quantify trabeculation in the 3-D image stacks (10 2-D images) with 20-fold improvement in efficiency as compared to our previous method. This approach allows for further quantification of the 3-D hypertrabeculation and ventricular remodeling in response to ventricular injury and regeneration.

Advances in imaging technologies are in parallel with the development and application of Saak transform. We have recently reported a sub-voxel light-sheet microscopy to provide high-throughput volumetric imaging of mesoscale specimens at the cellular resolution [54]. Integrating this method with Saak transform would further enhance spatiotemporal resolution and image contrast with a large field-of-view. The Saak transform would also be applicable to other molecular imaging modalities [55, 56], including optical coherence tomography, photoacoustic tomography and magnetic resonance imaging, for the post-image processing of in vivo imaging data set.

## V. CONCLUSION

Elucidating tissue injury and repair would accelerate the fields of developmental biology and regenerative medicine. We integrated Saak transform-based machine learning with LSFM imaging to assess cardiac architecture in zebrafish model in response to chemotherapy-induced injury, providing a minimal training data set for efficient and accurate image segmentation. The Saak transform-based LSFM, in conjunction with edge detection, allows for both quantitative and qualitative insights into cardiac ultrastructure and organ morphogenesis. Integrating the augmented kernels with random forest classifier captures the human’s inference capacity to enhance the accuracy of classification. Overall, our platform establishes the basis for quantitative computation of high throughput microscopy images to advance imaging and computational analysis for both fundamental and translational research.

## VII. ACKNOWLEDGMENT

All authors thank Dr. Xiaolei Xu at Mayo Clinic for providing adult zebrafish model of chemotherapy-induced injury, and appreciate Dr. René R Sevag Packard at UCLA for providing cardiac light-sheet images of adult zebrafish.

